# Effects of a rifampicin pre-treatment on linezolid pharmacokinetics

**DOI:** 10.1101/572610

**Authors:** Fumiyasu Okazaki, Yasuhiro Tsuji, Yoshihiro Seto, Chika Ogami, Yoshihiro Yamamoto, Hideto To

## Abstract

Linezolid is an oxazolidinone antibiotic that effectively treats methicillin-resistant *Staphylococcus aureus* (MRSA) and vancomycin-resistant *Enterococci* (VRE). Since rifampicin induces other antibiotic effects, it is combined with linezolid in therapeutic regimes. However, linezolid blood concentrations are reduced by this combination, which increases the risk of the emergence of antibiotic-resistant bacteria. We herein demonstrated that the combination of linezolid with rifampicin inhibited its absorption and promoted its elimination, but not through microsomal enzymes. Our results indicate that the combination of linezolid with rifampicin reduces linezolid blood concentrations via metabolic enzymes.

## Introduction

Linezolid is an oxazolidinone antibiotic that is used in the treatment of methicillin-resistant *Staphylococcus aureus* (MRSA) and vancomycin-resistant enterococci (VRE). Linezolid inhibits initiation complex formation of the 70S ribosome by binding to the 50S ribosomal subunit[1]. This mechanism of antibiotic action differs from those of other antibiotics, and, thus, linezolid is not cross-resistant to other antibiotics [2]. Therefore, these characteristics of linezolid are advantageous for decreasing the risk of antibiotic-resistant bacteria emerging due to reductions in blood linezolid concentrations when administered in combination with other drugs.

Rifampicin is an anti-tuberculosis drug that inhibits RNA polymerase β, and is effective against MRSA. The combination of rifampicin with one or more drugs is recommended in order to prevent the emergence of rifampicin-resistant bacteria. As rifampicin penetrates biofilms and exerts bactericidal antibacterial effects, it has been the treatment of choice for infection control [3]. However, rifampicin-resistant bacteria may emerge when rifampicin is administered alone. Therefore, the use of rifampicin in combination with other antibiotics is strongly recommended [4, 5].

The combination of rifampicin with linezolid has been shown to inhibit the generation of rifampicin-resistant bacteria [6], and also induces the effects of linezolid [7, 8]. However, numerous linezolid combination studies with rifampicin demonstrated that blood linezolid concentrations decreased when it was combined with rifampicin [9, 10]. Reductions in blood linezolid concentrations may increase the risk of drug treatment failure as well as the emergence of antibiotic-resistant bacteria. Therefore, the mechanisms underlying the pharmacokinetic interactions between linezolid and rifampicin need to be elucidated in more detail. The observed reduction of linezolid concentration when combined with rifampicin has gained attention recently. Although these reductions have been extensively examined, specific studies have not yet been performed.

Therefore, the present study had 2 aims: (1) to examine if rifampicin pre-treatment promotes the metabolism or elimination of linezolid in an *in vivo* study, and (3) to clarify whether the induction of cytochrome P450 (CYP) by a rifampicin pre-treatment promotes the degradation of linezolid in an *in vitro* study.

## Materials and Methods

### Animals

Five-week-old male ICR mice (Sankyo Labo Service Corporation, Inc., Tokyo, Japan) were housed under a standard light/dark cycle (light phase: 7:00-19:00) at a temperature of 24 ± 1°C and humidity of 60 ± 10% with *ad libitum* access to food and water. Anesthesia was maintained with intermittent administration of intraperitoneal pentobarbital. All experiments were performed in accordance with the Guide for the Care and Use of Laboratory Animals distributed by the U.S. National Institutes of Health [11] and the rules and regulations of animal experiments in University of Toyama (permit number: A2012PHA-45).

### RT-PCR analysis

Mice were orally administered rifampicin (100 mg/kg) or vehicle (control group) once per day for 7 days. One day after the final administration, livers were removed from mice. Total mRNA was extracted using RNAiso Plus (TaKaRa Bio Inc., Otsu, Japan) and cDNA was synthesized with the PrimeScript RT reagent Kit with the gDNA Eraser (TaKaRa Bio Inc.). Real-time PCR was performed using the KOD SYBR qPCR Mix (TOYOBO, Kita, Japan) with StepOnePlus (Thermo Fisher Scientific, Waltham, MA, USA). All samples were normalized with the housekeeping gene *Gapdh*.

### Collection of serum after the administration of linezolid

Mice were orally administered rifampicin (100 mg/kg) or vehicle (control group) once per day for 7 days. One day after the final rifampicin administration, linezolid (25 mg/kg) was orally or intraperitoneally administered to mice. Blood samples were obtained from the heart after the administration of linezolid. Whole blood was allowed to clot, and sera were collected by centrifugation at 3,000 ×*g* for 15 min.

### In vitro pharmacokinetics study

Mice were orally administered rifampicin (100 mg/kg) or vehicle (control group) for 7 days. Rifampicin (100 mg/kg) was orally administered to mice once per day for 7 days. One day after the final rifampicin administration, the liver was removed from mice. Total protein was extracted using T-PER Tissue Protein Extraction Reagent (Thermo Fisher Scientific). Protein concentrations were measured using the BCA Protein Assay Kit (Thermo Fisher Scientific). Linezolid solution (final concentration, 25 μg/mL) and NADPH (final concentration, 2.5 mM) were added to the total protein solution (final concentration, 20 μg/mL) and incubated at 37°C. Methanol was added to stop the reaction 0.5, 1, 2, and 4 h after the incubation. Linezolid concentrations were measured using high performance liquid chromatography (HPLC). Dexamethasone (final concentration, 12.5 μg/mL) was used as a positive control.

### Measurement of drug concentrations by HPLC

Linezolid concentrations were measured using an HPLC method with ultraviolet (UV) detection, according to a previously reported method [12]. Dexamethasone concentrations were measured using an HPLC method with UV detection. The HPLC system (Shimadzu Corporation) consisted of a LC-2010 pump, LC-2010 autosampler, LC-2010 UV detector, and LC-2010 column oven. Data were collected and analyzed using LC solution. Separation was performed on an ODS Hypersil column (Cadenza 5CD-C18, 150 mm × 4.6 mm, 5 µm; Imtakt Co.). A solution of 1% phosphoric acid was used for the mobile phase, and pH was adjusted to 5 by the addition of 10 M sodium hydroxide. The pump flow rate was 1.0 mL/min. The column temperature was maintained at 40°C. The wavelength of optimum UV detection was set at 254 nm. Calibration curves were linear over a concentration range of 1 to 100 µg/mL. Intra/inter-day CV was less than 5.0%, and LLOQ was 1 μg/mL for dexamethasone concentrations.

### Statistical analysis

An unpaired *t*-test was used to analyze differences between two groups. Dunnett’s test was used for post-hoc comparisons. Differences between the groups with a P value of < 0.05 were considered to be significant.

## Results

### Influence of the rifampicin pre-treatment on linezolid pharmacokinetics *in vivo*

In order to investigate whether a rifampicin pre-treatment influences the absorption, metabolism, and elimination of linezolid, mice were orally administered linezolid one day after the rifampicin pre-treatment and blood linezolid concentrations were measured using HPLC. The rifampicin pre-treatment reduced linezolid blood concentrations at all sampling points, and decreased AUC by 30% from that in the control group (Figure 1A). In order to clarify whether the rifampicin pre-treatment promotes the metabolic or eliminated process, linezolid was intravenously administered one day after an injection of rifampicin and linezolid blood concentrations were then measured using HPLC. The results obtained showed that linezolid blood concentrations were reduced by the rifampicin pre-treatment (Figure 1B).

**Figure 1.**
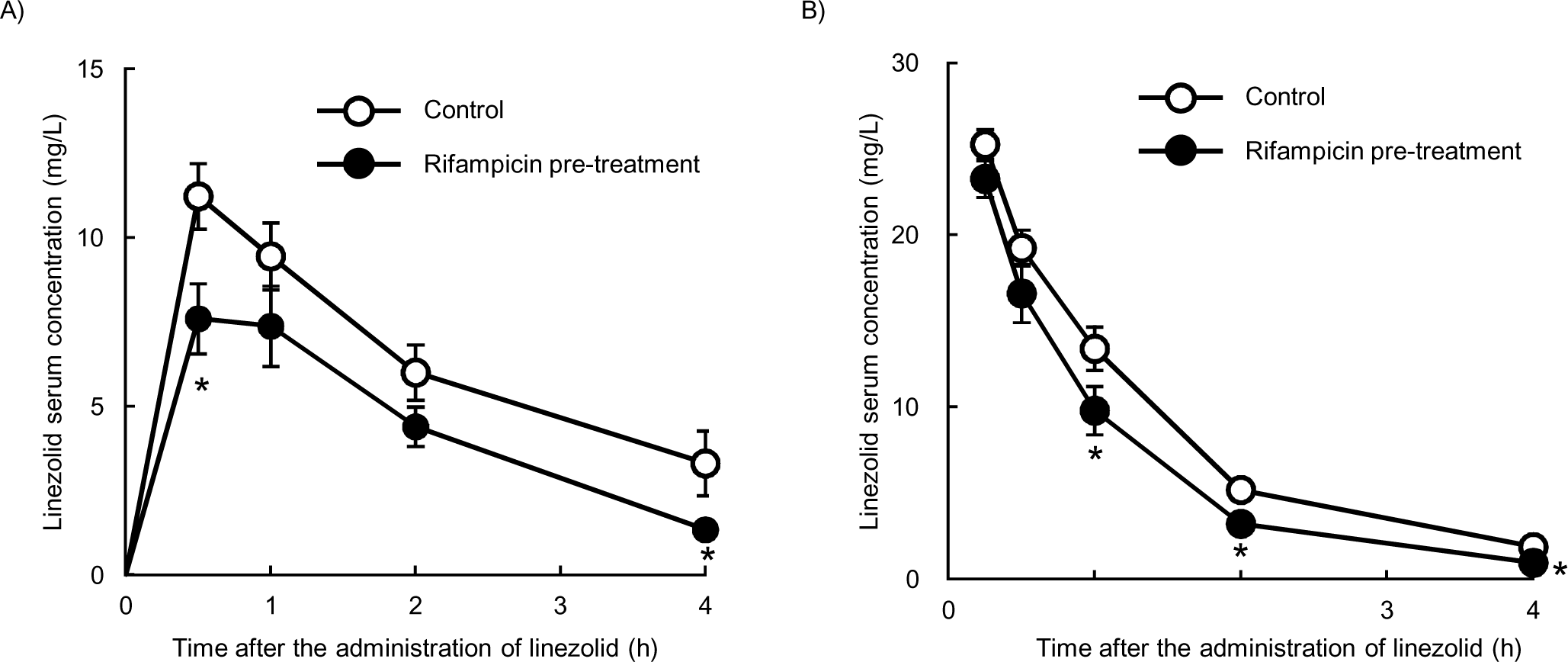
Effects of the rifampicin pre-treatment on the pharmacokinetics of linezolid. (A) Linezolid (25 mg/kg) was orally administered to mice (Control group). Mice were orally administered rifampicin (100 mg/kg) for 7 days. One day after the rifampicin pre-treatment, linezolid (25 mg/kg) was orally administered to mice (Rifampicin pre-treatment group). (B) Linezolid (25 mg/kg) was intravenously administered to mice (Control group). Mice were orally administered rifampicin (100 mg/kg) for 7 days. One day after the rifampicin pre-treatment, linezolid (25 mg/kg) was intravenously administered to mice (Rifampicin pre-treatment group). * *P* < 0.05 (unpaired *t*-test). Data are represented as the mean ± S.D. (n = 3 for each group).

### Influence of the rifampicin pre-treatment on the expression of *Cyp3a11* and drug degradation in liver metabolic enzymes

Rifampicin induces the expression of CYP3A4, and mouse *Cyp3a11* is homologous with human *CYP3A4* [13–15]. We investigated the induction of *Cyp3a11* expression by the rifampicin pre-treatment, and found that *Cyp3a11* levels were higher than those in the control group (Figure 2A): a 4-fold increase in the liver and a 40-fold increase in the intestines. Liver contain a number of enzymes that metabolize drugs, including CYPs. In order to investigate the reductions induced in linezolid concentrations by enzymes in liver metabolic enzymes, we measured linezolid concentrations in an *in vitro* study. Linezolid concentrations were not reduced by the extract solution by liver pre-treated with rifampicin. Since dexamethasone is metabolized by CYPs, we performed the same experiment using dexamethasone as a positive control. Dexamethasone concentrations were reduced by the extract solution by liver pre-treated with rifampicin (Figure 2B).

**Figure 2.**
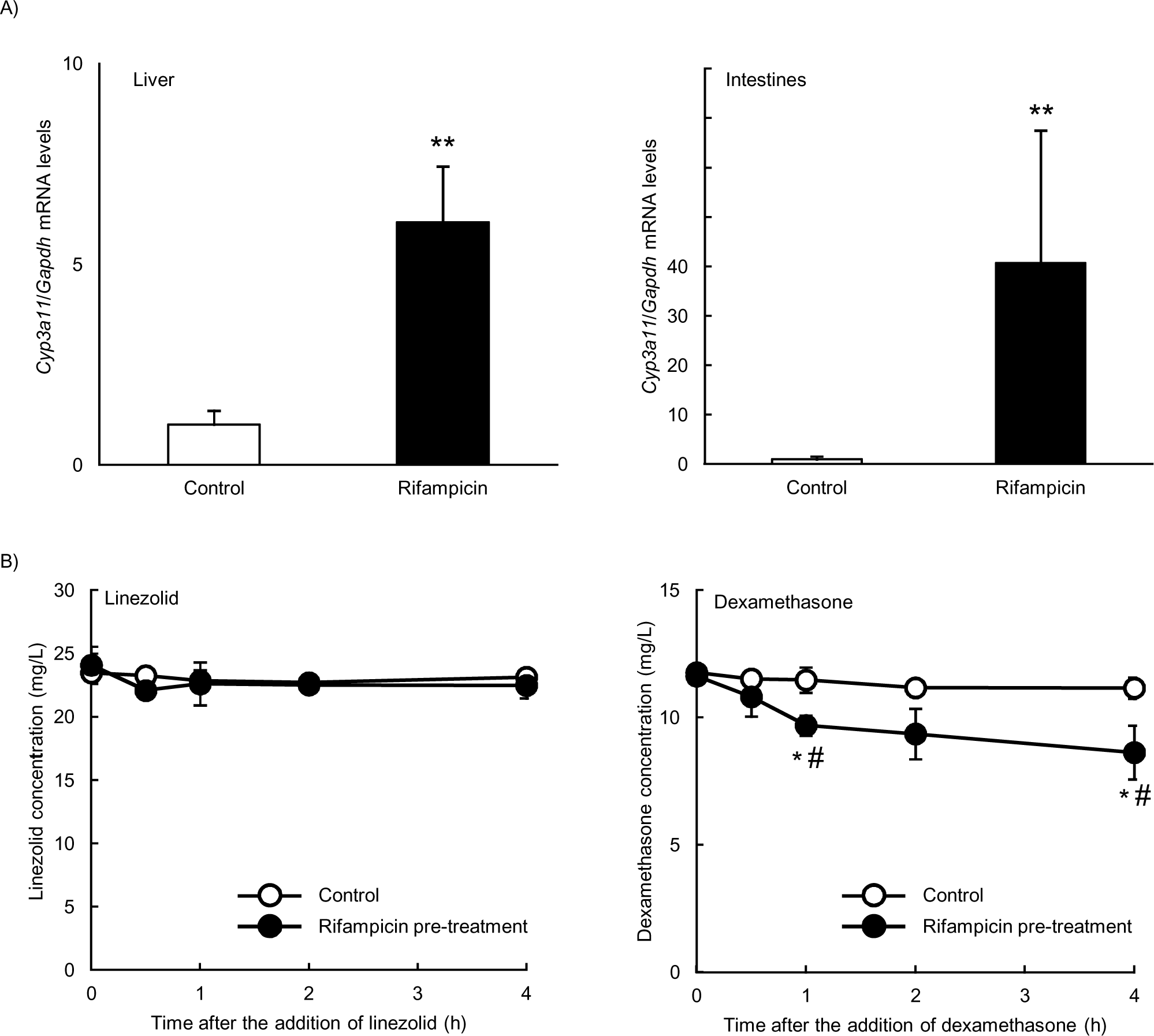
Influence of the induction of CYPs on the pharmacokinetics of linezolid. (A) Effects of the rifampicin pre-treatment on *Cyp3a11* expression. Mice were orally administered rifampicin (100 mg/kg) or vehicle (control group) for 7 days. ** *P* < 0.01 (unpaired *t*-test). (B) Effects of the rifampicin pre-treatment on linezolid and dexamethasone concentrations in liver metabolic enzymes. * *P* < 0.05 (unpaired *t*-test), # *P* < 0.05 vs 0 h (Dunnett’s test). Data are represented as the mean ± S.D. (n = 3 for each group).

## Discussion

Linezolid uses a novel mechanism of action against microbes, and its antibiotic effects are induced when it is combined with other antibiotics. Since linezolid is not metabolized by CYPs, it does not affect the pharmacokinetics of other antibiotics [2, 16]. However, linezolid blood concentrations are reduced when it is combined with rifampicin, and this has recently been attracting increasing attention [9, 17].

In the present study, in order to avoid direct interactions with rifampicin, mice were treated with rifampicin once per day for 7 days until one day before the administration of linezolid. Linezolid blood concentrations were reduced by the rifampicin pre-treatment. To the best of our knowledge, rifampicin is a substrate of some organic anion transporting polypeptides (OATPs), but does not induce the expression of these transporters. P-gp is an efflux transporter, and its inhibition has been shown to increase AUC, thereby reducing drug elimination [18]. Previous studies reported that rifampicin is a pregnane X receptor (PXR) ligand that induces P-gp [19, 20]. Rifampicin affects the concentrations of other drugs through P-gp [21]. However, linezolid is not a substrate of P-gp or OATP according to Pfizer, and a previous study also suggested that these transporters do not affect its pharmacokinetics [9]. Therefore, the present results indicated that efflux transporters did not affect the pharmacokinetics of linezolid after the rifampicin pre-treatment.

Rifampicin also induces CYP3A expression via PXR [22]. Dexamethasone is a substrate of CYPs, and its concentration decreased in the *in vitro* experiments conducted in the present study, whereas that of linezolid did not. Previous studies suggested that linezolid is not a substrate of or is poorly metabolized by CYPs [2, 16]. Moreover, the total protein solution, extracted from liver after rifampicin or dexamethasone treatment, were used in *in vitro* study. Therefore, these findings suggest that the pharmacokinetics of linezolid are influenced by factors without liver metabolic enzymes.

Undesired reductions in blood antibiotic concentrations lead to failed drug treatments and the emergence of antibiotic-resistant bacteria. We herein demonstrated that linezolid blood concentrations were reduced by the rifampicin pre-treatment, which affected drug absorption and metabolism/elimination. Liver metabolic enzymes did not affect the pharmacokinetics of linezolid with the rifampicin pre-treatment. Previous studies suggested that linezolid is metabolite by the lactone and lactam pathway [23, 24]. Although it is unclear that rifampicin induces the lactone and lactam pathway, these results suggest that transporters and liver metabolic enzymes poorly affect the pharmacokinetics of linezolid in rifampicin combination therapy.

## Funding

This work was supported by JSPS KAKENHI Grant Number 17K08438.

## Ethical approval

Approval was not required.

## Acknowledgments

This work was supported by the Life Science Research Center, University of Toyama.

## Conflict of interests

The authors declare no conflict of interest associated with this manuscript.

## References

1. Hamel JC, Stapert D, Moerman JK, Ford CW. Linezolid, critical characteristics. Infection. 2000;28(1):60-4. Epub 2001/02/07. PubMed PMID: 10744479.

2. Fung HB, Kirschenbaum HL, Ojofeitimi BO. Linezolid: an oxazolidinone antimicrobial agent. Clinical therapeutics. 2001;23(3):356–91. Epub 2001/04/25. PubMed PMID: 11318073.

3. Perlroth J, Kuo M, Tan J, Bayer AS, Miller LG. Adjunctive use of rifampin for the treatment of Staphylococcus aureus infections: a systematic review of the literature. Archives of internal medicine. 2008;168(8):805–19. Epub 2008/04/30. doi: 10.1001/archinte.168.8.805. PubMed PMID: 18443255.

4. Forrest GN, Tamura K. Rifampin combination therapy for nonmycobacterial infections. Clinical microbiology reviews. 2010;23(1):14–34. Epub 2010/01/13. doi: 10.1128/cmr.00034-09. PubMed PMID: 20065324; PubMed Central PMCID: PMCPMC2806656.

5. Sanders WE, Jr. Rifampin. Annals of internal medicine. 1976;85(1):82–6. Epub 1976/07/01. PubMed PMID: 937928.

6. Vergidis P, Rouse MS, Euba G, Karau MJ, Schmidt SM, Mandrekar JN, et al. Treatment with linezolid or vancomycin in combination with rifampin is effective in an animal model of methicillin-resistant Staphylococcus aureus foreign body osteomyelitis. Antimicrobial agents and chemotherapy. 2011;55(3):1182–6. Epub 2010/12/30. doi: 10.1128/aac.00740-10. PubMed PMID: 21189340; PubMed Central PMCID: PMCPMC3067063.

7. Cabellos C, Garrigos C, Taberner F, Force E, Pachon-Ibanez ME. Experimental study of the efficacy of linezolid alone and in combinations against experimental meningitis due to Staphylococcus aureus strains with decreased susceptibility to beta-lactams and glycopeptides. Journal of infection and chemotherapy: official journal of the Japan Society of Chemotherapy. 2014;20(9):563–8. Epub 2014/06/30. doi: 10.1016/j.jiac.2014.05.008. PubMed PMID: 24973908.

8. Drusano GL, Neely M, Van Guilder M, Schumitzky A, Brown D, Fikes S, et al. Analysis of combination drug therapy to develop regimens with shortened duration of treatment for tuberculosis. PloS one. 2014;9(7):e101311. Epub 2014/07/09. doi: 10.1371/journal.pone.0101311. PubMed PMID: 25003557; PubMed Central PMCID: PMCPMC4086932.

9. Gandelman K, Zhu T, Fahmi OA, Glue P, Lian K, Obach RS, et al. Unexpected effect of rifampin on the pharmacokinetics of linezolid: in silico and in vitro approaches to explain its mechanism. Journal of clinical pharmacology. 2011;51(2):229–36. Epub 2010/04/08. doi: 10.1177/0091270010366445. PubMed PMID: 20371736.

10. Ashizawa N, Tsuji Y, Kawago K, Higashi Y, Tashiro M, Nogami M, et al. Successful treatment of methicillin-resistant Staphylococcus aureus osteomyelitis with combination therapy using linezolid and rifampicin under therapeutic drug monitoring. Journal of infection and chemotherapy: official journal of the Japan Society of Chemotherapy. 2016;22(5):331–4. Epub 2016/01/07. doi: 10.1016/j.jiac.2015.11.012. PubMed PMID: 26732509.

11. National Research Council Committee for the Update of the Guide for the C, Use of Laboratory A. The National Academies Collection: Reports funded by National Institutes of Health. In: th, editor. Guide for the Care and Use of Laboratory Animals. Washington (DC): National Academies Press (US) National Academy of Sciences.; 2011.

12. Tsuji Y, Holford NHG, Kasai H, Ogami C, Heo YA, Higashi Y, et al. Population pharmacokinetics and pharmacodynamics of linezolid-induced thrombocytopenia in hospitalized patients. British journal of clinical pharmacology. 2017;83(8):1758–72. Epub 2017/02/12. doi: 10.1111/bcp.13262. PubMed PMID: 28186644; PubMed Central PMCID: PMCPMC5510085.

13. Staudinger JL, Goodwin B, Jones SA, Hawkins-Brown D, MacKenzie KI, LaTour A, et al. The nuclear receptor PXR is a lithocholic acid sensor that protects against liver toxicity. Proceedings of the National Academy of Sciences of the United States of America. 2001;98(6):3369–74. Epub 2001/03/15. doi: 10.1073/pnas.051551698. PubMed PMID: 11248085; PubMed Central PMCID: PMCPMC30660.

14. Bodin K, Lindbom U, Diczfalusy U. Novel pathways of bile acid metabolism involving CYP3A4. Biochimica et biophysica acta. 2005;1687(1-3):84–93. Epub 2005/02/15. doi: 10.1016/j.bbalip.2004.11.003. PubMed PMID: 15708356.

15. Holmstock N, Gonzalez FJ, Baes M, Annaert P, Augustijns P. PXR/CYP3A4-humanized mice for studying drug-drug interactions involving intestinal P-glycoprotein. Molecular pharmaceutics. 2013;10(3):1056–62. Epub 2013/01/31. doi: 10.1021/mp300512r. PubMed PMID: 23360470; PubMed Central PMCID: PMCPMC3594649.

16. Wynalda MA, Hauer MJ, Wienkers LC. Oxidation of the novel oxazolidinone antibiotic linezolid in human liver microsomes. Drug metabolism and disposition: the biological fate of chemicals. 2000;28(9):1014–7. Epub 2000/08/19. PubMed PMID: 10950842.

17. Egle H, Trittler R, Kummerer K, Lemmen SW. Linezolid and rifampin: Drug interaction contrary to expectations? Clinical pharmacology and therapeutics. 2005;77(5):451–3. Epub 2005/05/19. doi: 10.1016/j.clpt.2005.01.020. PubMed PMID: 15900290.

18. Mealey KL. Therapeutic implications of the MDR-1 gene. Journal of veterinary pharmacology and therapeutics. 2004;27(5):257–64. Epub 2004/10/27. doi: 10.1111/j.1365-2885.2004.00607.x. PubMed PMID: 15500562.

19. Tian R, Koyabu N, Morimoto S, Shoyama Y, Ohtani H, Sawada Y. Functional induction and de-induction of P-glycoprotein by St. John’s wort and its ingredients in a human colon adenocarcinoma cell line. Drug metabolism and disposition: the biological fate of chemicals. 2005;33(4):547–54. Epub 2005/01/11. doi: 10.1124/dmd.104.002485. PubMed PMID: 15640377.

20. Chan GN, Patel R, Cummins CL, Bendayan R. Induction of P-glycoprotein by antiretroviral drugs in human brain microvessel endothelial cells. Antimicrobial agents and chemotherapy. 2013;57(9):4481–8. Epub 2013/07/10. doi: 10.1128/aac.00486-13. PubMed PMID: 23836171; PubMed Central PMCID: PMCPMC3754350.

21. Ouwerkerk-Mahadevan S, Snoeys J, Peeters M, Beumont-Mauviel M, Simion A. Drug-Drug Interactions with the NS3/4A Protease Inhibitor Simeprevir. Clinical pharmacokinetics. 2016;55(2):197–208. Epub 2015/09/12. doi: 10.1007/s40262-015-0314-y. PubMed PMID: 26353895; PubMed Central PMCID: PMCPMC4756048.

22. MacLeod AK, McLaughlin LA, Henderson CJ, Wolf CR. Activation status of the pregnane X receptor influences vemurafenib availability in humanized mouse models. Cancer research. 2015;75(21):4573–81. Epub 2015/09/13. doi: 10.1158/0008-5472.can-15-1454. PubMed PMID: 26363009; PubMed Central PMCID: PMCPMC4634205.

23. MacGowan AP. Pharmacokinetic and pharmacodynamic profile of linezolid in healthy volunteers and patients with Gram-positive infections. The Journal of antimicrobial chemotherapy. 2003;51 Suppl 2:ii17-25. Epub 2003/05/06. doi: 10.1093/jac/dkg248. PubMed PMID: 12730139.

24. Slatter JG, Stalker DJ, Feenstra KL, Welshman IR, Bruss JB, Sams JP, et al. Pharmacokinetics, metabolism, and excretion of linezolid following an oral dose of [(14)C]linezolid to healthy human subjects. Drug metabolism and disposition: the biological fate of chemicals. 2001;29(8):1136–45. Epub 2001/07/17. PubMed PMID: 11454733.

